# Putting a New Spin of G4 Structure and Binding by Analytical Ultracentrifugation

**DOI:** 10.1101/359356

**Authors:** William L. Dean, Robert D. Gray, Lynn DeLeeuw, Robert C. Monsen, Jonathan B. Chaires

## Abstract

Analytical ultracentrifugation is a powerful biophysical tool that provides information about G-quadruplex structure, stability and binding reactivity. This chapter provides a simplified explanation of the method, along with examples of how it can be used to characterize G4 formation and to monitor small-molecule binding.

## 1. Introduction

Analytical ultracentrifugation (AUC) is underappreciated by the G-quadruplex (G4) community. AUC is a venerable biophysical technique that has a (nearly) one-hundred year history. Theodor Svedberg invented the analytical ultracentrifuge in 1925, and won the Noble Prize in Chemistry the next year for his research on colloids and protein using his invention. AUC has since been widely used as a fundamental technique for the determination of macromolecular structure, reaction stoichiometry and ligand affinity [1-4]. AUC is based on first-principle physical theory, and can be used to determine absolute molecular weights of molecules, along with their hydrodynamic shapes. Our laboratory has found AUC useful for a variety of G4 structural studies [5-13]. The intent of this chapter is to provide a simplified overview of AUC and then to show its utility for characterizing G4 structure and binding.

Figure 1 shows a schematic of the most basic AUC experiment, sedimentation velocity (SV). A sample is placed in one sector of centerpiece within a sealed cell assembly with quartz windows (Figure 1A). A reference solution is placed in the second sector. The cell assembly is then placed in a rotor and spun in an ultracentrifuge to produce a high centrifugal field. The centrifugal field is sufficient to cause the sedimentation of molecules within the sample cell in the direction of the field, toward the bottom of sector. The AUC instrument uses an optical system, typically absorbance, to scan the sample cell to monitor the concentration of molecules at each radial position in the cell, and to record their movement as a function of time at constant centrifugal force. As molecules move within the sample cell, a boundary is formed that changes with time. Figure 1B shows a schematic of an SV experiment, with intermittent scans showing the migration of a boundary toward the bottom of the sample cell. As the boundary migrates to the bottom it also broadens because of diffusion of the sedimenting molecule. Such primary data contain sufficient information to extract the structural properties of the sedimenting molecules.

**Figure 1.**
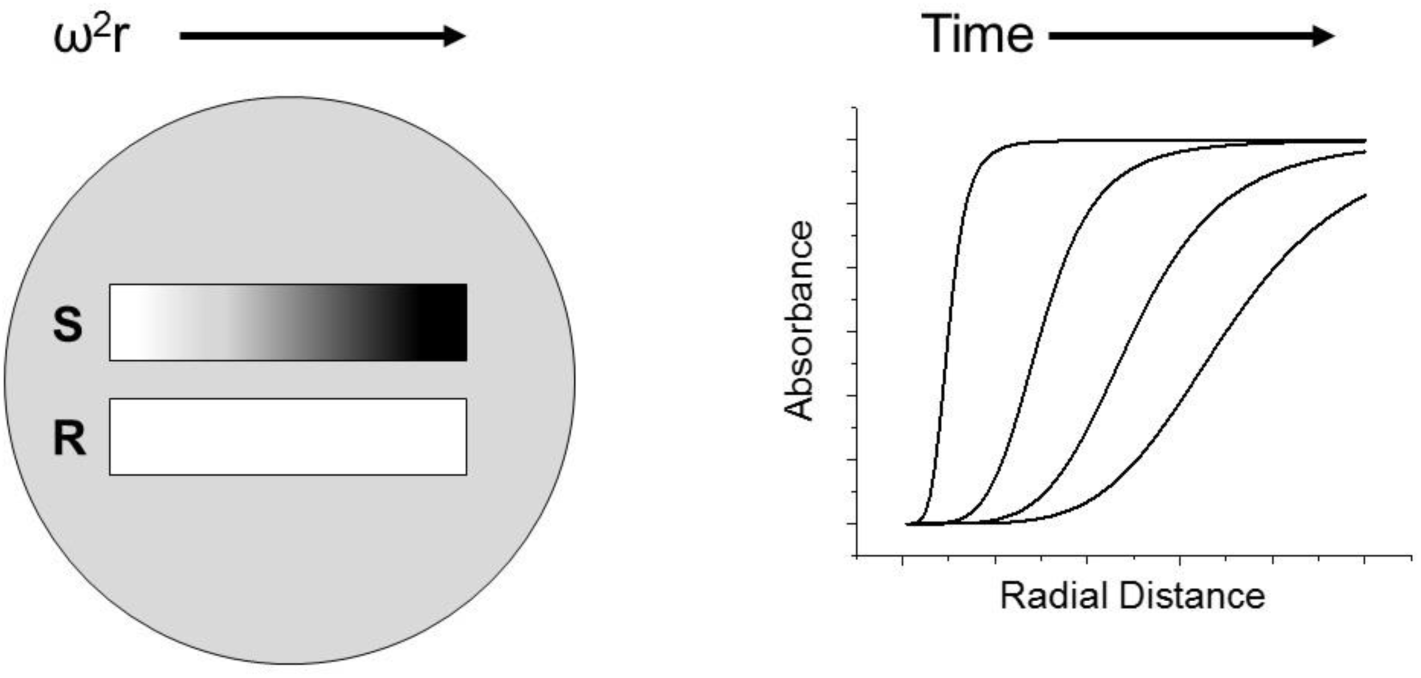
Schematic representation of a sedimentation velocity experiment. The panel on the left represents a double sector centrifuge cell with a sedimenting sample in the upper sector and the reference buffer in the lower sector. The right panel shows four absorbance scans taken at four different times during sedimentation.

As any number of basic textbooks derive and show, the sedimentation coefficient (*s*) is defined as the velocity (*v*) of the moving boundary (determined at the midpoint) divided by the centrifugal field strength (*ω^2^r*):

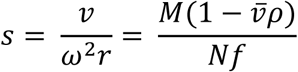

The sedimentation coefficient is then equal to product of the molecular weight of the molecule (*M*) and the buoyancy term 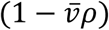 divided by the product of Avagadro’s number (N) and the frictional coefficient (*f*). In the buoyancy term, 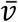 is the partial specific volume and ρ is the density of the solvent. The sedimentation coefficient is thus determined by the mass of the molecule and its shape. A useful extension of this basic equation is the *Svedberg equation*:

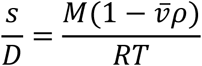

where *D* is the diffusion coefficient of the molecule, *R* the gas constant and *T* the absolute temperature. The molecular weight thus can be obtained from experimentally measured sedimentation and diffusion coefficients.

Mass spectrometry shares with AUC the ability to determine absolute molecular weights of G4 molecules. Both methods can be used to evaluate sample homogeneity or to determine strand stoichiometry and ligand binding. AUC, however, offers some unique advantages. First, AUC measurements are done in solutions of defined and invariant composition, and sedimentation is sensitive to not only the mass of molecules but also their hydrodynamic shape. Second, as molecules are separated in mass spectrometry the equilibrium of the initial sample solution is necessarily perturbed, perhaps leading to some strand dissociation in multi-stranded G4 structures or small-molecule ligand dissociation. In contrast, in AUC measurements the moving G4 boundary remains in equilibrium as it moves through the cell, minimizing perturbation of multi-stranded structures or of bound ligand.

## 2. Materials

Oligonucleotides were from IDT, Coralville, IA or Eurofins Genomics, Louisville, KY. Stock solutions were prepared by dissolving the lyophilized, desalted oligos in MQ-H_2_0 to a final concentration of ~1 mM. Working samples were prepared by diluting the stock DNA to ~3-5 μM in the appropriate buffer. In sample preparation, the annealing protocol used can play an important role in quadruplex formation [14]. All samples except those at pH 11.5 were annealed by heating for 5-10 min in a 1-liter boiling water bath followed by slow (12–24 h) cooling to room temperature.

## 3. Methods

### 3.1 Analytical ultracentrifugation

Operation of the analytical ultracentrifuge does not lend itself to simple step-by-step protocols. Mastery of the method requires some practice, and is best learned by the study of an excellent tutorial available at: http://www.analyticalultracentrifugation.com/sedphat/experimental_protocols.htm We have presented detailed protocols for the analysis of AUC data obtained for G4 structures in a previous volume of this series [13] and elsewhere [6].

### 3.2 Illustrative Results

Figure 2 shows an example of experimental data obtained by sedimentation velocity of the “hybrid” G4 structure formed by the human telomere repeat sequence Tel22, 5’AGGG(TTAGGG)_3_. With modern AUC instrumentation, digital data are collected and advanced software packages like SEDFIT or UltraScan can be used to fit the data in sophisticated ways to obtain s and D coefficients simultaneously, thereby providing both molecular weight estimates and shape information from a single sedimentation experiment. Figure 2B shows the results of such an analysis, presented as a c(s) plot in which the best-fit distribution of sedimentation coefficients derived from the data is shown. The results show that this G4 sample is homogeneous with a sedimentation coefficient of 1.9S.

**Figure 2.**
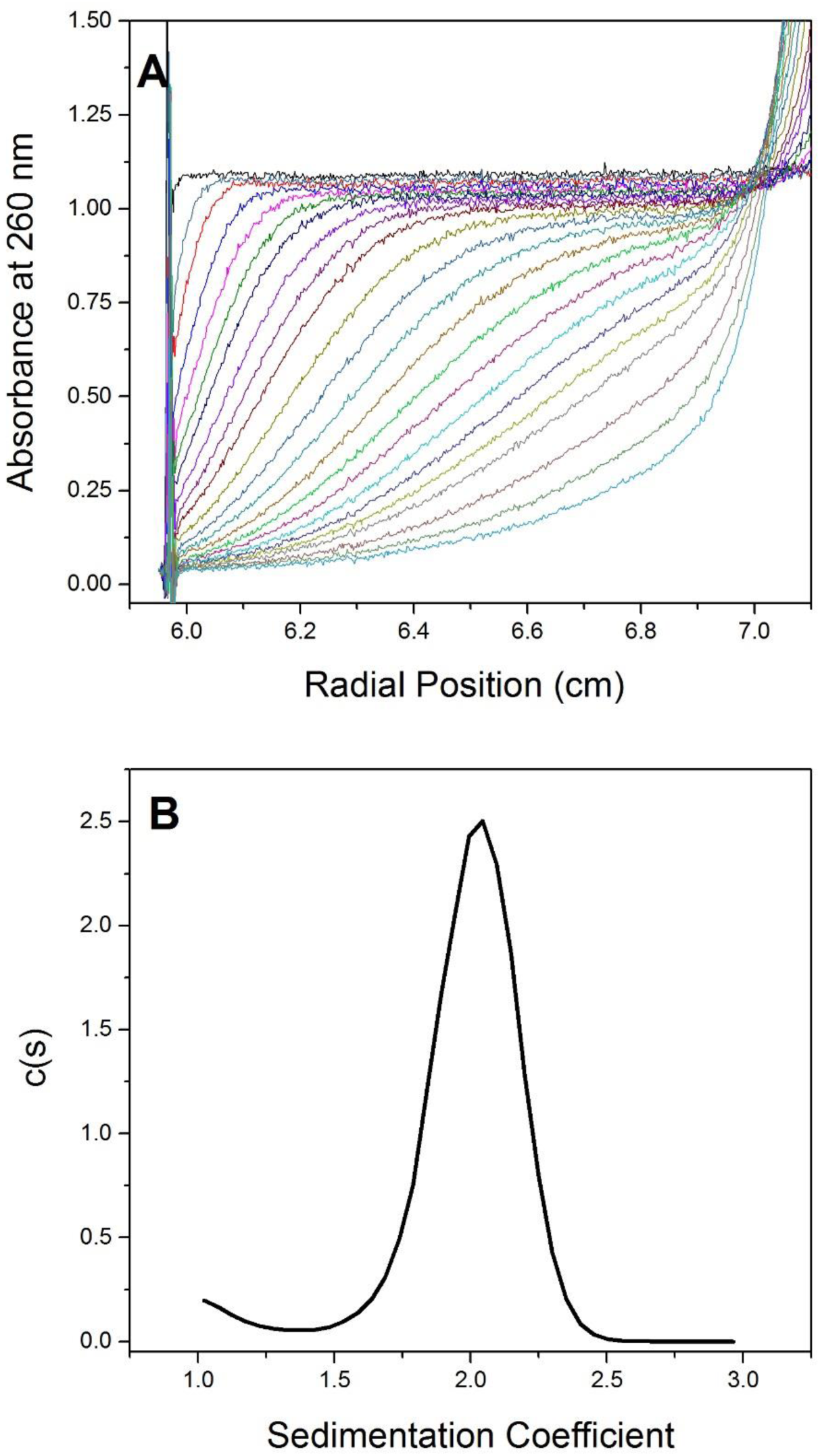
Sedimentation velocity experiment showing results from a solution containing 5 µM Tel22 in 0.2 M KCl. Panel A shows the raw data from 23 of the 100 scans taken over a 6 hour time period at 40,000 rpm and 20°C. Panel B shows sedfit analysis using the continuous c(s) distribution model. The results show that this G4 sample is homogeneous with a sedimentation coefficient of 1.9S.

### 3.3 Application of SV to demonstrate G4 structural formation

Figure 3 shows how SV can be used to demonstrate folding of an oligonucleotide into a compact structure. The telomeric sequence d[TTGGG(TTAGGG)_3_A] has been shown to fold into a compact 3 tetrad hybrid structure termed 2GKU in the presence of K^+^ [15]. We used AUC to show the different sedimentation behaviors of the unfolded sequence in LiCl, the unfolded sequence at pH 11.5, the folded 2GKU sequence in K^+^, an unstructured sequence of similar molecular weight, dT_24_, and finally the duplex formed with its complementary sequence. The differences in sedimentation behavior reflect differences in structure, not molecular weight, except for the case of the duplex. As indicated by the equation for the sedimentation coefficient above, an increase in the frictional coefficient due to unfolding will decrease the observed sedimentation coefficient as shown for 2GKU in Li_+_, at pH 11.5 and for the dT_24_ sequence. The fact that 2GKU in Li^+^ has a larger sedimentation coefficient than the sequence at pH 11.5 or dT_24_ suggests that some folding has been retained in Li^+^. The sedimentation coefficient of the duplex is increased due to the near doubling of molecular weight.

**Figure 3.**
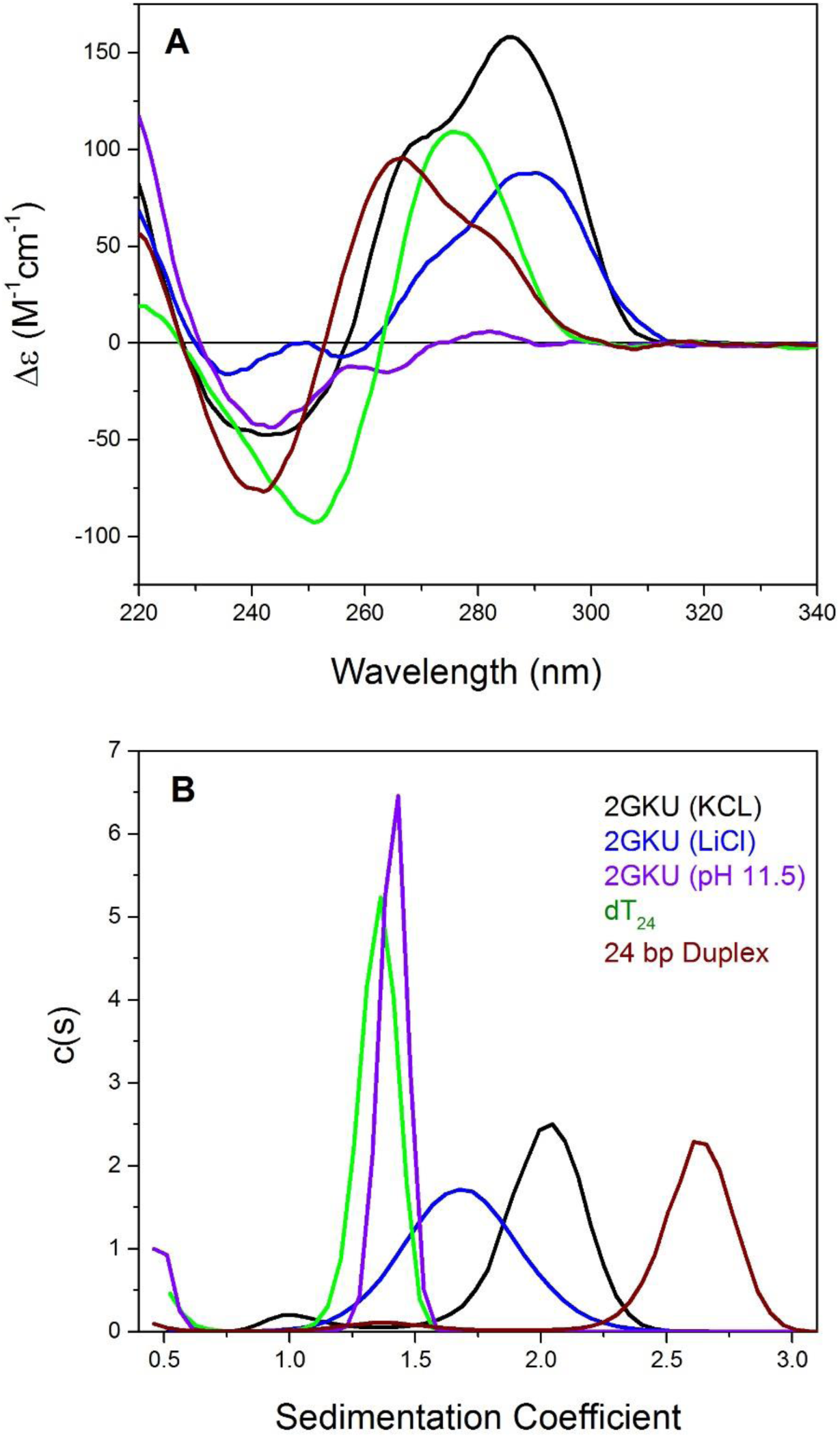
AUC and CD analysis of 2GKU under various buffer conditions known to result in the formation of different structures. Panel A shows CD spectra of 2GKU in KCl (black), 2GKU in LiCl (blue), 2GKU duplex (red), dT_24_ (green) and 2GKU at pH 11.5 (purple). The buffer was 10 mM sodium phosphate with added 185 mM KCl or LiCl, pH 7.0 or 11.5. Blank-corrected CD spectra were recorded in a 1-cm pathlength quartz cuvette at 25 °C with a Jasco J810 spectropolarimeter equipped with Pelteir temperature controller as described [30]. CD spectra were normalized using the equation Δɛ = θ/32980.l.c where is the observed ellipticity in millidegrees, l is the pathlength in cm and c is the molar strand concentration of DNA. Figure 3B shows sedfit analysis of sedimentation velocity results using the continuous c(s) distribution model of the same solutions using the same color scheme as in 3A.

The structures of 2GKU under the buffer conditions used for AUC were analyzed by circular dichroism as shown in Figure 3A. The conclusions with respect to oligonucleotide folding are consistent with the CD spectra presented in Figure 3A. For example, 2GKU in 185 mM KCl is fully folded into a hybrid-1 structure, while 2GKU in 180 mM LiCl gives a CD spectrum that indicates partial folding, probably a mixture of folded hybrid and unfolded forms. This is consistent with the known inability of Li^+^ to promote full G4 folding at these concentrations, but is inconsistent with the prevalent notion that Li^+^ is *completely* unable to promote G4 folding. As expected, 2GKU in 185 mM KCl, pH 11.5, is unfolded as indicated by CD. The GC-rich 2GKU duplex at neutral pH gives a CD spectrum characteristic of A-type DNA [16] while the CD spectrum of dT_24_ reflects the inherent asymmetry of the nucleobase chromophore itself in an non-helical (random) backbone conformation.

### 3.4 Detecting and characterizing G4 heterogeneity

A distinct and powerful advantage of SV is its ability to reveal and characterize heterogeneity. Figure 4 shows a case where a truncated form of the human telomere repeat sequence (5’(TTAGGG)_3_) was studied. It was anticipated that this sequence would fold into a triple-helical structure that might represent an intermediate along the multistate G4 folding pathway. SV shows, however, that this is not the case. There are two structures present at sedimentation coefficients of approximately 1.25 and 2.0 S and calculated molecular weights indicating a mixture of nearly equal amounts monomer and dimer. The CD spectrum (Figure 4 insert) is consistent with a mixture of conformations rather than that of a single structure. This result is expected based on the NMR studies of truncated Tel22 sequences [17].

**Figure 4.**
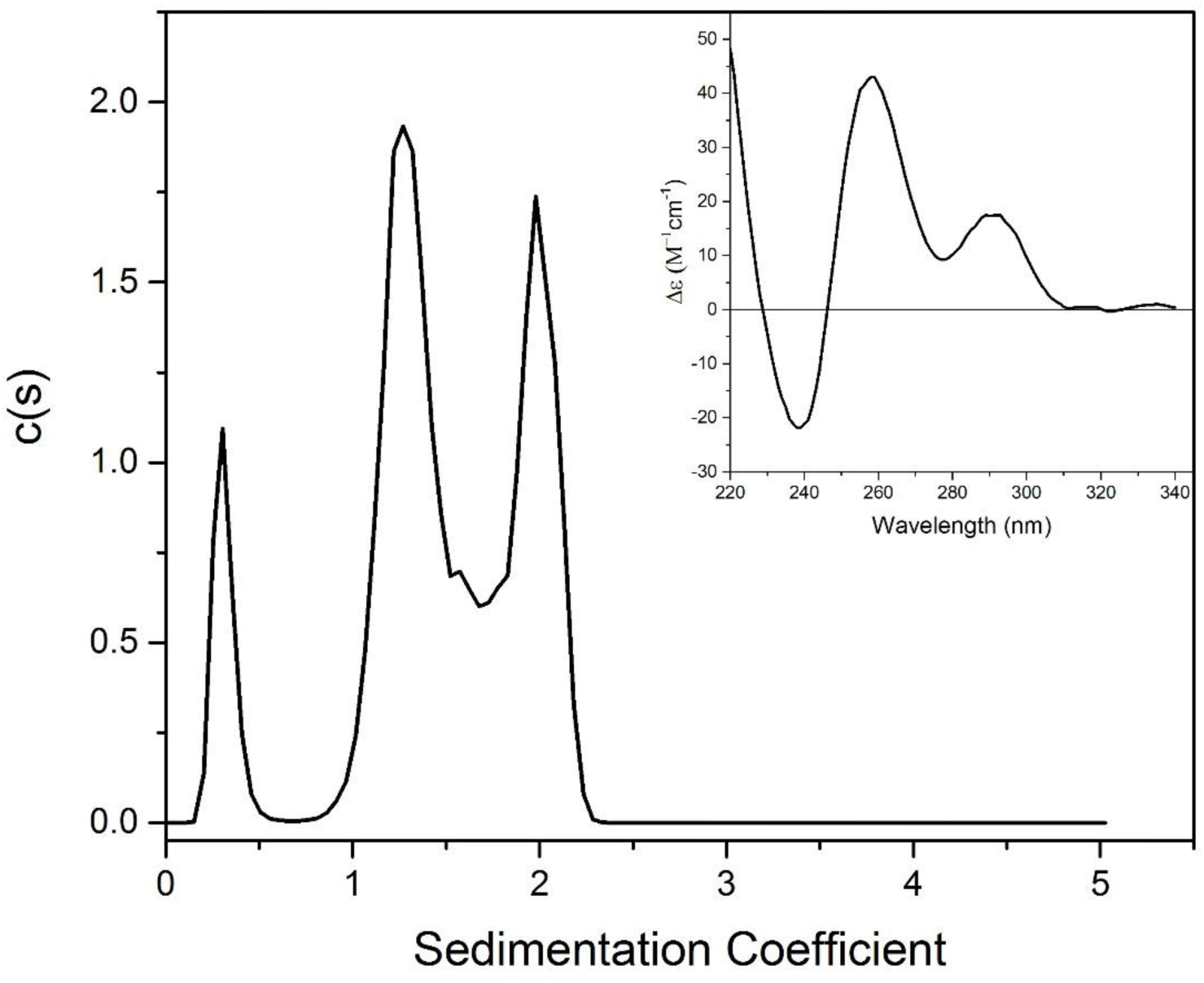
Sedfit analysis of 5’(TTAGGG)_3_ (7 μM) in 10 mM MOPS, 30 mM KCl, pH 7.0, using the continuous c(s) distribution model. Inset to Figure 4.CD spectrum of Tel22-5’Δ4 (7 μM) in 10 mM MOPS, 30 mM KCl, pH 7.0.

### 3.5 Application of SV to show ligand and protein binding to G4 structures

#### 3.5.1 General comments and considerations

Since the seminal paper by Steinberg and Schachman that described the use of the AUC absorbance optical system for studies of small-molecule binding to proteins [18], there have been myriad examples where AUC has been used to determine binding stoichiometry and dissociation constants for protein binding interactions. However, the use of AUC to study interactions between small molecules and DNA is highly underutilized, even though it was first used in 1954 (!) to show that methyl green binds to calf thymus DNA [19]. There are a few published examples utilizing AUC to study small molecule interactions with DNA [20,21,14], and even fewer assessing interaction of small molecules with G quadruplex-containing sequences [14,22].

The use of AUC to study small molecule-macromolecule interactions is based on first principles of thermodynamics. The determination of free and bound ligand using SV is actually quite straightforward if the ligand can be detected by absorbance, fluorescence or interference optical systems. Macromolecule and ligand are mixed and allowed to come to equilibrium. The resulting solution is then centrifuged in the AUC. After centrifugation for long enough to separate the faster sedimenting macromolecule from the much slower sedimenting ligand, the free concentration of the ligand can be determined directly from the portion of solution in the centrifuge cell that is free of macromolecule. Knowing the total amount of starting ligand, the amount bound ligand is calculated by difference. An idealized example is shown in Figure 5. This measurement provides the fractional saturation of the ligand at the total ligand concentration used in the experiment and a single point in a binding isotherm that can be used to determine the dissociation constant for the ligand. In addition one can obtain the stoichiometry of binding if the macromolecule is saturated, and an indication of any conformational change in the macromolecule that has occurred upon ligand binding based on changes in the sedimentation coefficient and frictional ratio. This methodology is analogous to equilibrium dialysis and to the classic Hummel and Dryer gel filtration chromatography binding method [23]. A major advantage of AUC over these other methods is that it allows for the analysis of systems free in solution without the possible complications of interactions with a matrix that can occur in gel filtration or the dialysis membrane. Furthermore, one also obtains structural information about the macromolecule in the presence of bound ligand.

**Figure 5.**
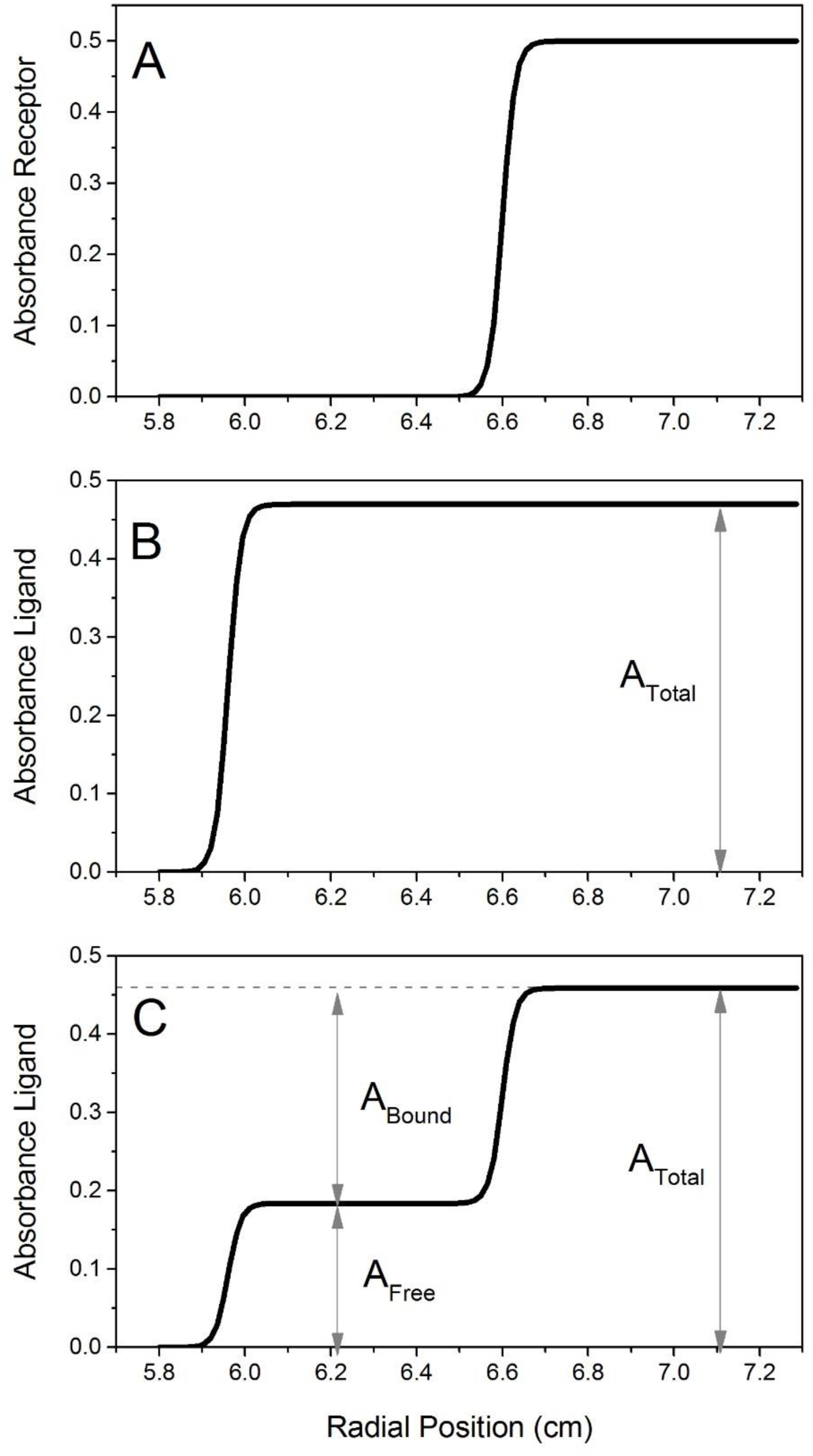
Idealized binding analysis experiment using sedimentation velocity with absorption optics. Panel A shows the results of a single scan at the wavelength where the receptor absorbs and at a time when the receptor has sedimented approximately 60% of the length of the solution column. Panel B shows the results at the same time at a wavelength where the ligand absorbs. Panel C shows analysis of the scan from panel B with concentrations of total, free and bound ligand clearly delineated.

If there is overlap between the absorption spectra of the ligand and macromolecule, then a multi-wavelength approach can be used to determine the amounts of ligand and macromolecule in the region of the cell containing both ligand and macromolecule [24-27]. For a two component system such as a single macromolecule and ligand, only two wavelengths are required to determine the contribution of macromolecule and ligand to each position in the centrifuge cell. New methodology has been developed for the most recent iteration of the analytical ultracentrifuge (Beckman Coulter Optima AUC) in which a multi-wavelength scan is acquired at each position in the cell during centrifugation [28]. A general methodology is outlined below.

#### 3.5.2 Steps to determine binding using absorbance optics in the Proteome Lab XL-A

1. The concentration of ligand should be sufficient to give an absorbance of 0.2-1.2.
2. If there is overlap between the absorbance of the ligand and the macromolecule, the experiment must be carried out at two wavelengths, preferably at the absorption maxima of the ligand and the macromolecule. The total absorbance of the two components at each wavelength should be in the range of 0.2-1.2.
3. 430 µL of sample is placed in the sample sector and 450 µL of buffer in the reference sector of a standard 2 sector cell used for sedimentation velocity.
4. A speed of 40,000 rpm is appropriate for most applications with G quadruplex-containing DNA oligonucleotides.
5. Buffers should contain at least 0.1 M salt to minimize non-ideal behavior.
6. To obtain s_20,w_ values and molecular weights, the density and viscosity of the buffer must be known. For simple buffers such as PBS, these parameters can be calculated using the program Sedinterp (free software: http://jphilo.mailway.com/download.htm).
7. Temperature can be maintained between 4 and 40° C in the AUC, but since DNA is quite stable, the standard temperature of 20° C is preferable.
8. Three cells are required for each binding experiment, one containing ligand only at the same concentration as used in the binding experiment, one cell with macromolecule alone at the same concentration used in the mixture, and 1 cell with a mixture of the two components.
9. After centrifugation sufficient for 100 scans (usually 4-8 hours), the data are analyzed by SEDFIT software (www.analyticalultracentrifugation.com) using the continuous c(s) distribution model. The analysis provides the starting concentration of each c(s) vs. s peak so that the concentration of the molecule in the peak can be directly calculated from the extinction coefficient at that wavelength.
10. The starting concentration absorbance for the small molecule is used to determine the free ligand concentration using the extinction coefficient and knowing the total small molecule concentration allows for calculation of bound ligand by difference.

#### 3.5.3 Analysis

Let L_T_ = total added ligand concentration, L_f_ = concentration of free ligand, L_b_ = concentration of bound ligand and M = total added macromolecule concentration in the cell containing the mixture of components. Also let A° _L_ = the absorbance of the ligand in the absence of macromolecule and ε_Lf_ = the extinction coefficient of the free ligand so that L_T_ = A°_L_/ε_Lf_∙l where l is the pathlength of the centrifuge cell (1.2 cm).

A_Lf_ = the absorbance of the free ligand after sedimentation of the mixture so that L_b_ = L_T_

– L_f_ = A°_L_/ε_Lf_∙l – A_Lf_/ε_Lf_∙l.

Then θ, fractional saturation = L_b_/L_f_+L_b_ and 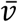(moles bound ligand/mole of macromolecule) = L_b_/M.

#### 3.5.4 Illustrative Results

.An example of a binding experiment in which interaction between a chemotherapeutic drug, Sunitinib and Tel48 (TTAGGG)_8_ is shown in Figure 6. At 418 nm only the drug absorbs, so the analysis is not complicated by the presence of Tel48 and looks similar to the idealized example in Figure 5 A. The concentration of free Sunitinib can be calculated directly from the absorbance at 418 nm after sedfit analysis in the continuous c(s) distribution model in which the area under the c(s) vs s peak is equal to the absorbance of that species (Figure 6B). Note that the absorbance for free Sunitinib from a single scan in Figure 6A is equal to the absorbance in the integrated peak in Figure 6B. In this experiment the starting concentration of Sunitinib was 30 µM and Tel48 was 3 µM, so at this 10-fold excess of drug, the apparent stoichiometry is 6.3 moles of drug/mole Tel48.

**Figure 6.**
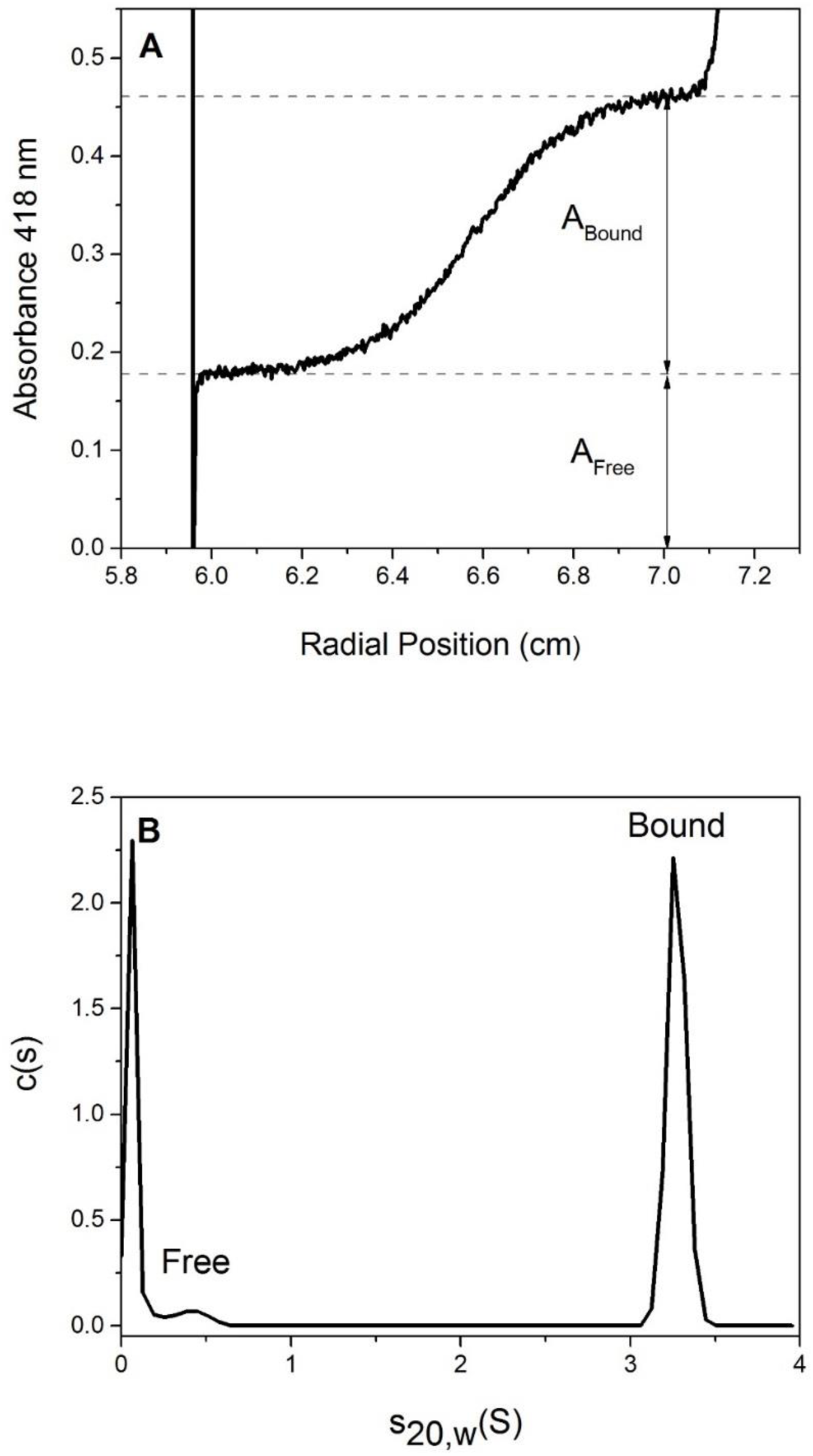
Sedimentation velocity analysis of binding of Sunitinib to Tel48. Sunitinib (30 µM) was mixed with Tel48 (3 µM) for 30 minutes and then analyzed by sedimentation velocity at 418 nm, the absorbance maximum of Sunitinib. Tel48 does not absorb at that wavelength. Panel A shows binding analysis as described in Figure 5C for a single scan where Tel48 has sedimented 60% of the length of the solution column. Panel B shows sedfit analysis of the binding experiment using the continuous c(s) distribution model. The area under the curve at the sedimentation coefficient of Tel48 represents bound Sunitinib and is exactly equal to the absorbance of bound Sunitinib determined in 6A, 0.28 absorbance units.

Table 1 summarizes the results for a number of small molecules found to interact with different G4 structures, including an antiparallel hybrid form (Tel 22), a structure containing contiguous antiparallel G4 structures (Tel48) and a parallel, “propeller” G4 structure (1XAV). The reliable determination of the binding stoichiometries show in Table 1 provides a firm foundation for the analysis of binding isotherms obtained by independent spectrophotometric titrations or by isothermal calorimetry.

**Table 1.** Summary of Small Molecule Binding AUC measurements in K+ buffers

| **Compound** | **DNA** | **Stoichiometry\*** | **Wavelength** |
| --- | --- | --- | --- |
| Sunitinib | Tel22 | 3 | 415 |
|  | Tel48 | 12 | 415 |
|  | 1XAV | 5 | 415 |
| Doxorubicin | Tel48 | 12 | 470 |
| Berberine | Tel22 | 1 | 345 |
|  | Tel48 | 2 | 345 |
| Ceritinib | Tel22 | 5 | 310 |
| Osimertinib | Tel22 | 3 | 310 |
|  | Tel48 | 4 | 320 |
| Ponatinib | Tel22 | 3 | 310 |
|  | Tel48 | 7 | 320 |
| Thiazole orange | Tel22 | 3 | 478 |
|  | 1XAV | 3 | 478 |
| NMM | Tel22 | 0 | 380 |
|  | 1XAV | 1 | 380 |
| Quindoline | Tel22 | 4 | 340 |
|  | 1XAV | 4 | 340 |
| Crizotinib | Tel22 | 2 | 315 |
|  | Tel48 | 6 | 315 |
| Bosutinib | Tel22 | 0 | 340 |
|  | Tel48 | 1 | 340 |
| Afatinib | Tel22 | 1 | 346 |
|  | Tel48 | 4 | 346 |
| Capecitabine | Tel22 | 1 | 300 |
|  | Tel48 | 2 | 300 |
| Palbociclib | Tel22 | 1 | 350 |
|  | Tel48 | 4 | 350 |

\*Rounded to nearest whole number. Ratio of small molecule:DNA for all experiments was a minimum of 10.

## 4. Final Comments

Our laboratory now regards AUC as an essential criterion for the demonstration of G4 formation in nucleic acid sequences of interest. AUC shows definitively whether or not a given sequence has folded into a compact structure and then provides the molecular weight of the structure to unambiguously determine its molecularity. AUC provides a clear characterization of the homogeneity of the folded structure, and clearly identifies the presence and exact amounts any unfolded species or unwanted aggregates. AUC clearly indicates if additional sample purification is needed. Once AUC has established that a sequence has folded into a homogeneous structure, additional methods like circular dichroism can be used with confidence to characterize the topological features of the folded G4 structure [29]. Our confidence in AUC as a biophysical tool based on fundamental physical principles is such that we have adopted the motto “*AUC non mentior*” – AUC does not lie.

## Acknowledgement

This work was supported by NIH Grant GM077422 and by the James Graham Brown Foundation.

